# Transcranial Focused Ultrasound to V5 Enhances Human Visual Motion Brain-Computer Interface by Modulating Feature-Based Attention

**DOI:** 10.1101/2023.09.04.556252

**Authors:** Joshua Kosnoff, Kai Yu, Chang Liu, Bin He

**Author notes:** Correspondence: Bin He, PhD.

## Abstract

Paralysis affects roughly 1 in 50 Americans. While there is no cure for the condition, brain-computer interfaces (BCI) can allow users to control a device with their mind, bypassing the paralyzed region. Non-invasive BCIs still have high error rates, which is hypothesized to be reduced with concurrent targeted neuromodulation. This study examines whether transcranial focused ultrasound (tFUS) modulation can improve BCI outcomes, and what the underlying mechanism of action might be through high-density electroencephalography (EEG)-based source imaging (ESI) analyses. V5-targeted tFUS significantly reduced the error for the BCI speller task. ESI analyses showed significantly increased theta activity in the tFUS condition at both V5 and downstream the dorsal visual processing pathway. Correlation analysis indicates that the dorsal processing pathway connection was preserved during tFUS stimulation, whereas extraneous connections were severed. These results suggest that V5-targeted tFUS’ mechanism of action is to raise the brain’s feature-based attention to visual motion.

## 1. INTRODUCTION

Paralysis affects roughly 1 in 50 Americans^1^. Over 70% of these people are younger than 65 years old, and over 40% of them reported unable to work. Unfortunately, there is no cure for paralysis. However, symptoms may be managed using brain-computer interfaces (BCIs), which decode brain signals to carry out a task, thereby bypassing the paralyzed regions. BCIs can read brain signals through either invasively implanted arrays or through non-invasive techniques like electroencephalography (EEG). Invasive BCIs have demonstrated potential to control a robotic arm^2–5^, decode speech^4,6,7^, and decode handwriting^8^, but the required brain surgery to implant the arrays severely limit their use cases. As a result, non-invasive BCIs present a more feasible option for widespread application. Non-invasive BCIs have demonstrated ability to move a robotic arm^9,10^, control a wheelchair^11^, or type on a virtual keyboard^12,13^, but even recent innovations in non-invasive BCIs have online accuracy rates of approximately 70 to 80%^9,14–16^, which leave room for improvement before seeking widespread clinical use.

A potential way to improve BCI is by pairing them with non-invasive neuromodulation devices. Traditional devices, such as transcranial electric stimulation and transcranial magnetic stimulation, have limited spatial specificity, on the orders of centimeters^17^, which modulate not only the targeted brain areas, but also those off-target^18^. This, combined with the challenge in temporal resolution, lead to these devices being used before or in-between BCI sessions to facilitate cortical learning, instead of during the sessions^19–21^. Perhaps for these reasons, behavioral results for these device modulated BCIs are mixed^19–21^. Transcranial focused ultrasound (tFUS), however, is an emerging technology that has been shown to have lateral precision on the order of millimeters^22^.

Previous research has demonstrated the capability of tFUS to modulate cortical circuits^22–25^ *during* the stimulation sessions^22–24^. Additional studies have also highlighted its ability to penetrate deep brain structures^26^ and activate microglia to clear Aβ plaque^27^. It has even been used in brain-brain interfaces where information was read from one subject’s brain via EEG and corresponding tFUS-evoked perceptions were elicited in another subject^28^. However, to date, it has not been used to modulate the brain signals that directly control BCI. In the present study, we set out to examine tFUS neuromodulation effects on improving BCI control through human behavior and noninvasive electrophysiological recordings.

The BCI paradigm tested is a motion-onset visual evoked potential (mVEP) BCI speller, which uses moving lines across a virtual keyboard to induce visual motion-based event related potentials (ERPs)^12,15,29,30^. The driving ERP of this paradigm is believed to be the N200, which is a negative deflection that peaks 150 to 250 ms after stimulus onset, though the N100 (a negative deflection roughly 100 ms post-stimulus)^31^ and the P300 (a positive deflection approximately 300 ms post-stimulus)^13^ are also associated with visual processing and BCI spellers. The primary brain area associated with visual motion processing is V5, or the middle temporal complex^32,33^. A prior tFUS study has demonstrated sonication of this area leads to improved visual motion detection^24^.

Our study explores, for the first time, whether tFUS can be used to improve BCI control by modulating V5 during an mVEP BCI speller task. We perform neural data analysis at both the EEG sensor level and the brain region-specific source domain to examine possible mechanisms of action.

## 2. RESULTS

### tFUS to V5 significantly reduced mVEP BCI speller error

The Euclidean errors for each letter typed by each subject were calculated by measuring the spatial distance between the target letter and the typed letter, with further normalizations with respect to the subject-specific epoch scans and ultrasound dosage (see Methods for details). A Kruskal-Wallis test indicated significant differences in medians across conditions (*p* < 0.01). Specific condition-to-condition comparisons were made with a Wilcoxon ranked sum test and Bonferroni-Holm *p* adjustment (Fig. 1). Results indicated that the Euclidean error for the tFUS condition (*N* = 21 subjects / 331 trials; error = 8.64 ± 11.8%) was significantly lower than those of the non-modulated (*N* = 21 subjects / 317 trials; error = 11.7 ± 12.8%; *p*_adjusted_ < 0.01) and decoupled-sham conditions (*N* = 15 subjects / 229 trials; error = 13.8 ± 16.9%; *p*_adjusted_ < 0.05). There was no significant difference detected between the decoupled-sham and non-modulated condition (*p*_adjusted_ > 0.05).

**Figure 1.**
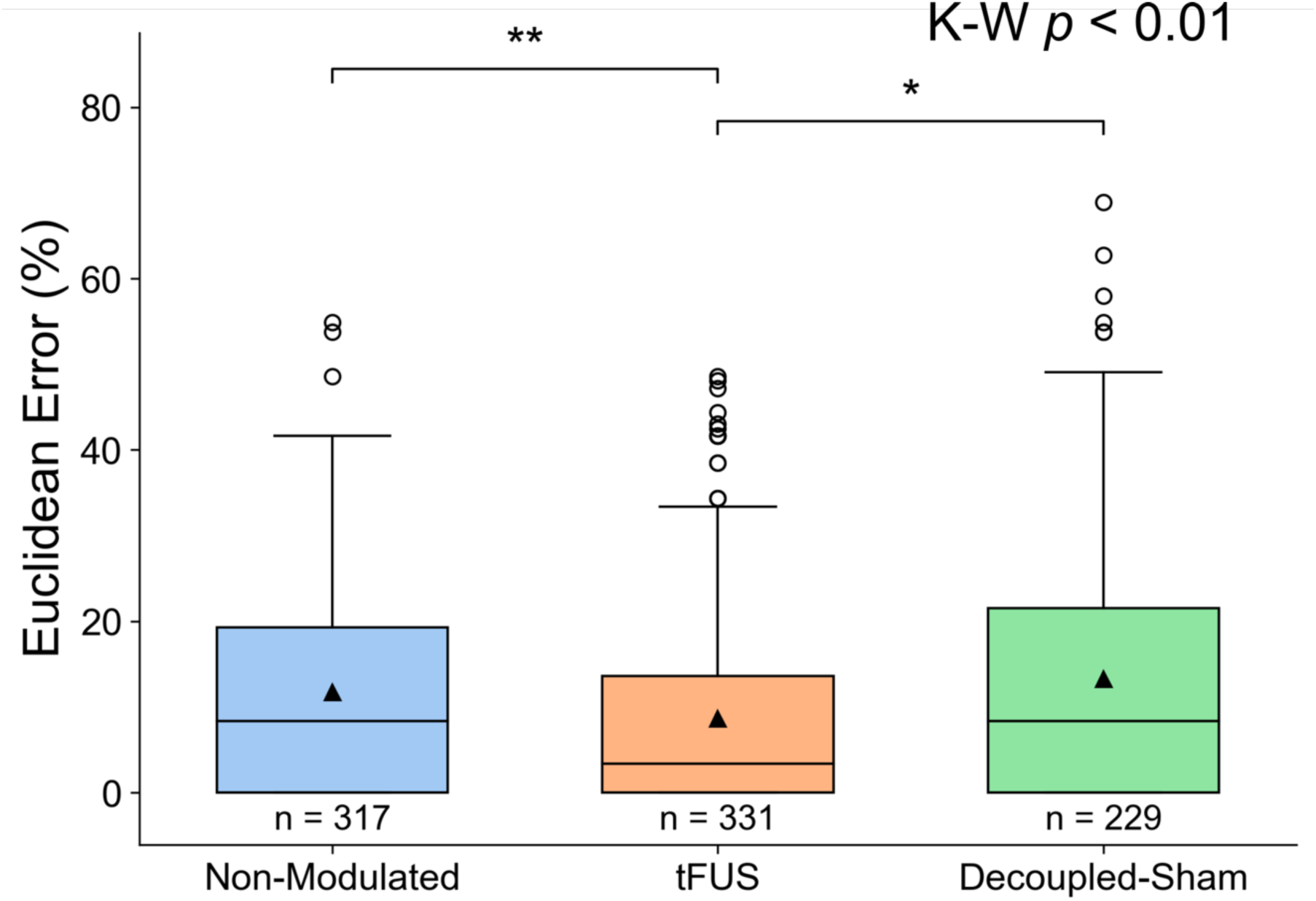
TFUS to V5 significantly improves mVEP BCI speller outcomes. A Kruskal-Wallis test indicated significant differences (*p* < 0.01) between the median Euclidean errors for each condition. tFUS sonication to V5 (*N* trials = 331 trials; mean error = 8.64 ± 11.8%) leads to significantly lower Euclidean error for the mVEP BCI speller compared to non-modulated (*N* trials = 317; mean error = 11.7 ± 12.8%) and decoupled-sham (*N* trials = 229; mean error = 13.8 ± 16.9%) conditions. No significant difference between the non-modulated and the decoupled-sham condition was found. Error was normalized with respect to the square root of the scans per trial and number of tFUS sonication per trial (Equation 1). Two-tailed Wilcoxon ranked sum test (with Bonferroni-Holm *p* adjustment) key: \**p*_adjusted_ < 0.05, \*\**p*_adjusted_ < 0.01.

### tFUS to V5 significantly alters detected EEG sensor-level signals

Preprocessed EEG sensor data were compared across tFUS (*N* subjects = 21 / trials = 1,364), decoupled-sham (*N* subjects = 15 / trials = 756) and non-modulated (*N* subjects = 20 / trials = 1,417 trials) conditions using MNE Python’s non-parametric permutation cluster F-test. The outcome revealed significant spatiotemporal clusters (*p* < 0.05) between tFUS and the control conditions (decoupled-sham: Fig. 2c top; non-modulated: Fig. 2c bottom) in the occipital regions between 140 and 340 ms post stimulus onset. The time-frequency cluster tests yielded significant differences across all three conditions in electrode *PO3* (Fig. 2d, left) in 3 to 9 Hz responses between 130 and 170 ms post stimulus onset (Fig. 2d, right).

**Figure 2.**
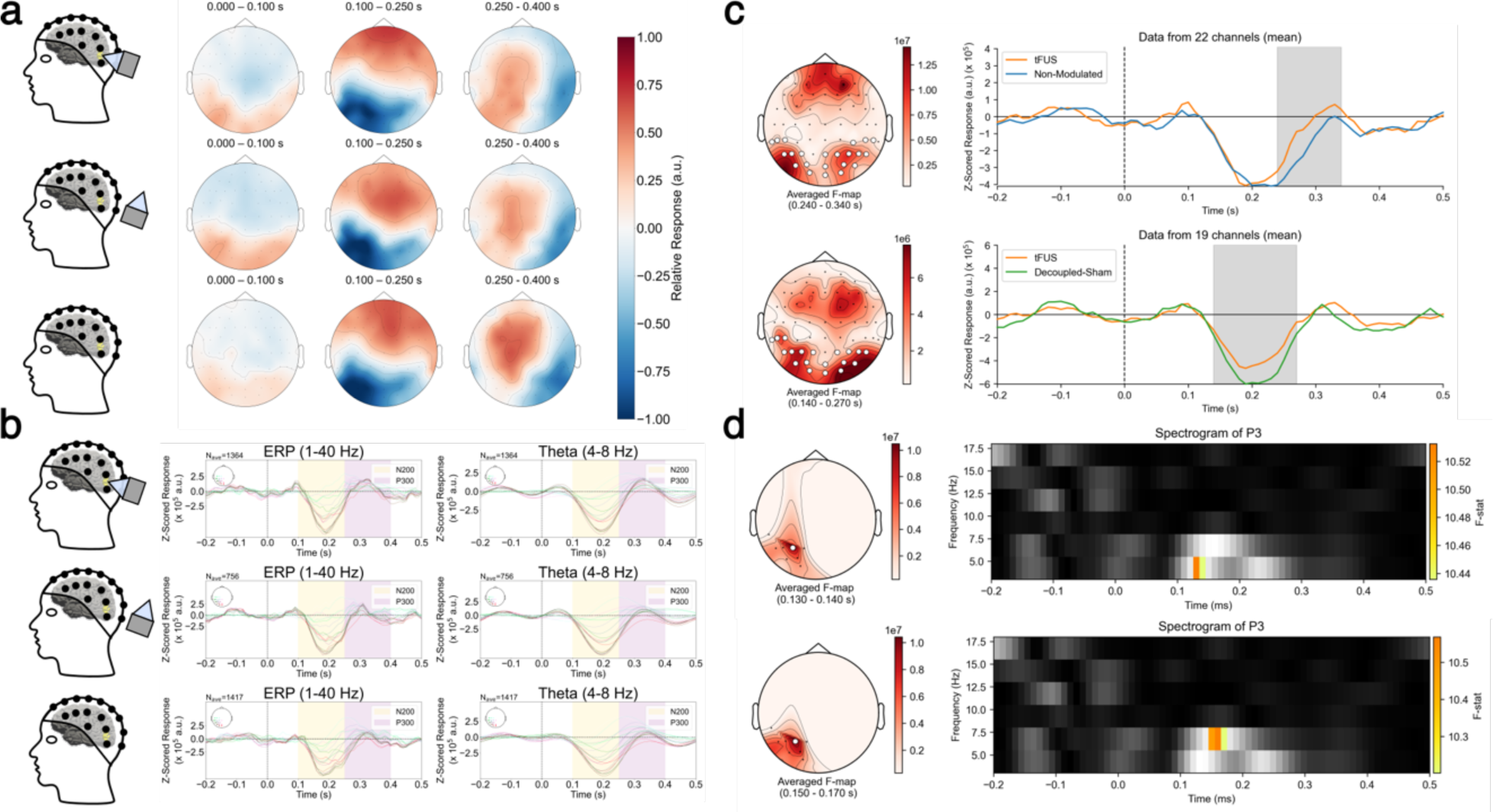
Significant differences are found between conditions in the EEG sensor domain. a) Topographic maps of the trial’s averaged activity for 0 to 100 ms (left), 100 to 250 ms (middle) and 250 to 400 ms (right) post stimulus filtered between 1 to 40 Hz for tFUS (top), decoupled-sham (middle), and non-modulated (bottom) conditions. The topo colormap is scaled to the relative max/min response for each condition’s z-score. b) Averaged left posterior electrode responses for 1 to 40 Hz response (left) and theta response (right) for tFUS (top), decoupled-sham (middle) and non-modulated condition (bottom). The approximate N200 and P300 waveform responses are highlighted in yellow and purple as the 100 to 250 ms and 250 to 400 ms windows, respectively. c) Significant spatiotemporal cluster (*p* < 0.05) between conditions using MNE Python’s nonparametric spatiotemporal cluster test with 1000 permutations when comparing tFUS against non-modulated (top) and decoupled-sham (bottom) conditions. (left) The F-statistics of a significant spatial cluster denoted by white circles. (right) The mean activity across channels and trials of the three trial conditions. Gray shaded regions indicate a significant temporal cluster corresponding to the spatial cluster. d) Significant spatiotemporal-frequency clusters when comparing all three conditions using MNE Python’s nonparametric spatiotemporal cluster test with 1000 permutations. (left) The F-statistic of electrode cluster (averaged F statistic threshold > 10). (right) The F-stat for significant time-frequency clusters (*p* < 0.05).

### tFUS to V5 significantly amplifies V5 N200 theta band power

To take the analysis a step further, EEG data were back-projected to the brain source domain by means of electrophysiological source imaging (ESI)^34^ using FreeSurfer^35,36^ and MNE Python^37^ and z-scored with respect to the 200 ms pre-stimulus baseline. N200 theta powers were calculated by time-frequency Morlet waveform power transformation of the 100 to 250 ms post stimulus window (Fig. 3b). A Kruskal-Wallis test considering the effect of ultrasound among three conditions yielded a significant difference in medians of N200 powers (*p* < 0.0001). Inter-condition comparisons indicated tFUS (6.98 ± 5.49 a.u.) to V5 during mVEP BCI significantly amplifies the N200 theta response in the V5 region compared to the non-modulated (5.57 ± 4.16 a.u.; *p*_adjusted_ < 0.0001) and decoupled-sham (4.99 ± 3.60 a.u.; *p*_adjusted_ < 0.0001) conditions. The decoupled-sham condition was also found to be significantly damped (*p*_adjusted_ < 0.05) compared to the non-modulated condition.

**Figure 3.**
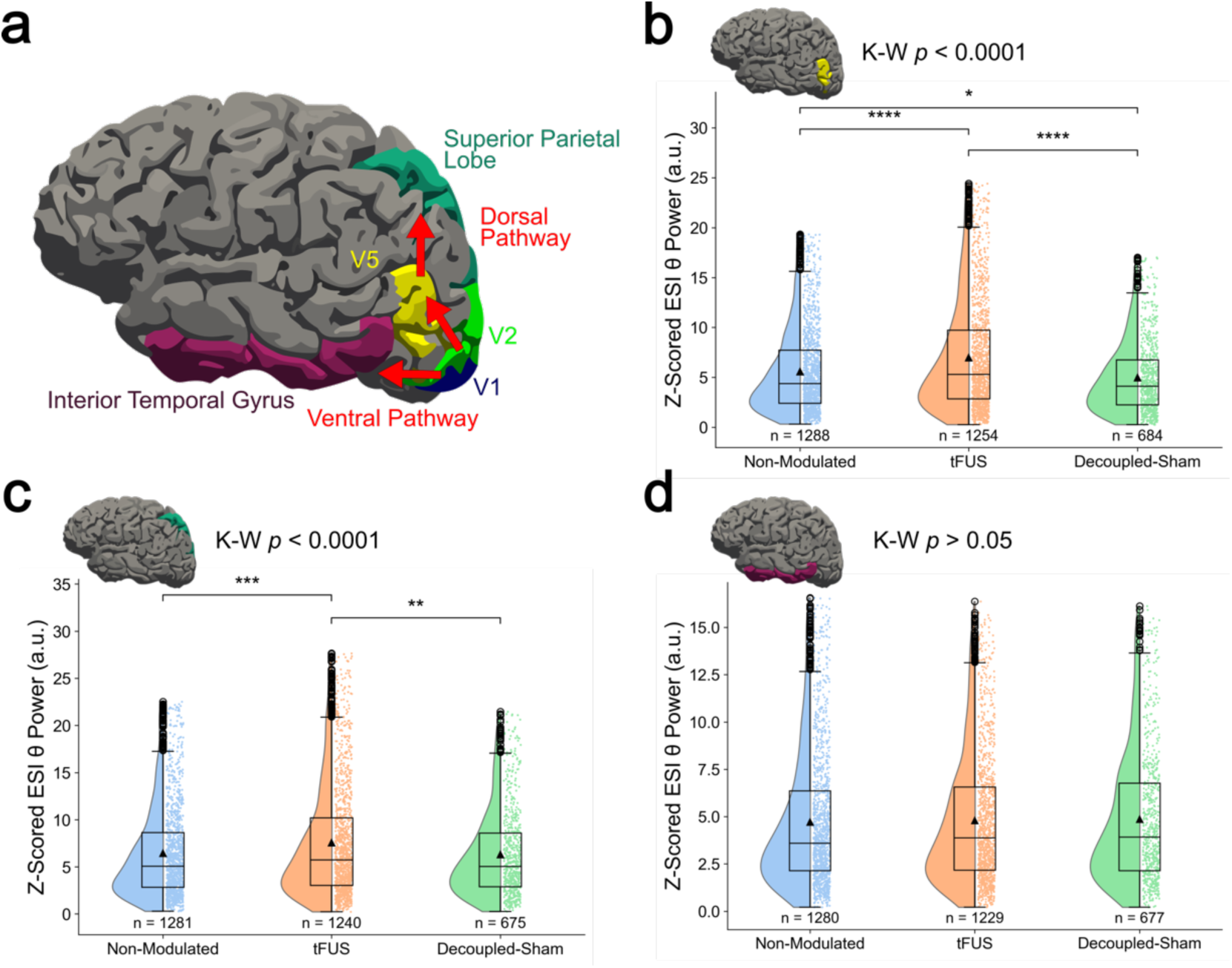
V5 activity is carried through the dorsal pathway. a) The FreeSurfer labeled regions for the cortical regions of the ventral and dorsal visual processing pathways. b) EEG source imaging of V5 yields a significant amplification of the N200 theta power for tFUS modulation over decoupled-sham and non-modulated conditions, as well as significant damping of the decoupled-sham compared to the non-modulated condition. c) EEG source imaging of the superior parietal lobe shows significant N200 theta amplification in the tFUS condition compared with the non-modulated and decoupled-sham. d) No significant difference is detected for the inferior temporal gyrus source image location. A Kruskal-Wallis test indicated significant differences in medians of amplitudes across conditions for V5 and superior parietal source imaged sites (b-c). Two-tailed Wilcoxon ranked sum test key with Bonferroni-Holm adjustment: * *p*_adjusted_ < 0.05, ** *p*_adjusted_ < 0.01, *** *p*_adjusted_ < 0.001, **** *p*_adjusted_ < 0.0001.

### tFUS to V5 theta-band amplification is continued downstream in the dorsal pathway

N200 theta power analysis was repeated for the superior parietal lobe (Fig. 3b) and the inferior temporal lobe (Fig. 3c), areas commonly associated with the dorsal and ventral visual processing pathways, respectively^38–40^. Kruskal-Wallis testing indicated significant differences in the median power across all three conditions (*p* < 0.0001) in the superior parietal lobe. Wilcoxon ranked sum test results for the superior parietal lobe showed significant amplification of N200 power for the tFUS condition (7.57 ± 6.01 a.u.) over non-modulated (6.44 ± 4.88 a.u.; *p*_adjusted_ < 0.001) and decoupled-sham (6.32 ± 4.62 a.u.; *p*_adjusted_ < 0.01). No significant difference was found between the non-modulated and decoupled-sham conditions (*p*_adjusted_ > 0.05). When examining the ventral pathway (inferior temporal gyrus), no significant difference was found among the conditions (Kruskal-Wallis *p* > 0.05; all *p*_adjusted_ > 0.05; tFUS: 4.80 ± 3.46 a.u.; decoupled-sham: 4.87 ± 3.51 a.u.; non-modulated: 4.71 ± 3.56 a.u.).

### Modulation of BCI behavioral results and theta band EEG are dependent on the tFUS targeting location

The effects of steering the ultrasound away from V5, but still projecting ultrasound energy to the brain, were also investigated (Fig. 4). Comparing this “US-control” condition (*N_testing_* = 12 subjects / 172 BCI letter trials; *N_training_* = 13 subjects / 873 ESI scan trials; mean Euclidean error = 12.4 ± 13.5%) to V5-targeted tFUS resulted in significant differences for both BCI errors (*p* < 0.01) and theta powers in V5 (US-Control: 6.30 ± 5.13 a.u.; *p* < 0.01) and the superior parietal lobe (US-Control: 6.53 ± 4.99 a.u.; *p* < 0.01). A significant difference was also found in the inferior temporal gyrus theta powers (US-Control: 4.29 ± 3.07 a.u.; *p* < 0.01).

**Figure 4.**
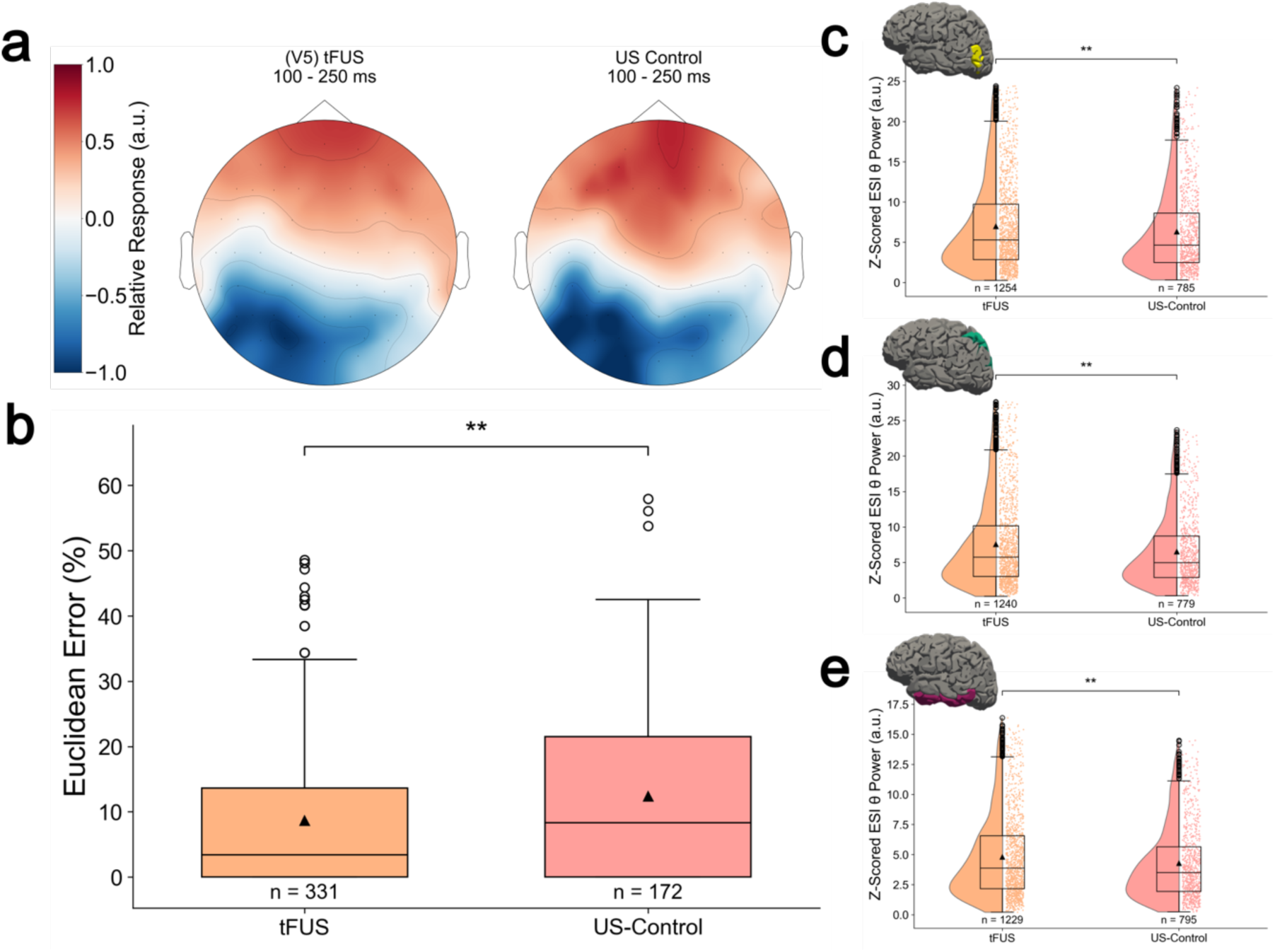
Behavioral results and EEG source imaging data are significantly different for tFUS to V5 compared to control location. a) Trial averaged topographic maps for tFUS to V5 and ultrasound control location from 100 to 250 ms post stimulus, filtered between 1 to 40 Hz. The colormap is scaled to the relative max/min of each condition’s z-score. b) There was a significant increase in Euclidean error for the ultrasound control location (12.4 ± 13.5%) compared to V5- targeted tFUS (8.64 ± 11.8%) c) The V5 source imaged N200 theta power was significantly increased for tFUS to V5 condition compared to the control location. d) Superior parietal source imaging results reflect the same significant difference as the V5 location. e) Theta power of the interior temporal gyrus is significantly damped in the US-Control condition compared to V5- targeted tFUS. Results were analyzed with a two tailed Wilcoxon ranked sum test. \*\**p* < 0.01.

### Correlation analysis reveals tFUS to V5 breaks extraneous connections and maintains task-relevant ones

Cortical connections were quantified using Pearson’s correlation coefficient for the non-modulated (Fig. 5a) and tFUS (Fig. 5b) conditions. The correlation coefficients for each subject were pooled together and tested for significant differences using a Wilcoxon ranked sum test. Results indicate that the correlation between V5-IT is significantly lowered (*p* < 0.05; Fig. 5d) when tFUS is applied to V5, whereas the information flow through the dorsal pathway (V5-SP) is not significantly altered (*p* > 0.05; Fig. 5c). There is also a significant (*p* < 0.05) disconnection of the V1-SP pathway for the tFUS condition.

**Figure 5.**
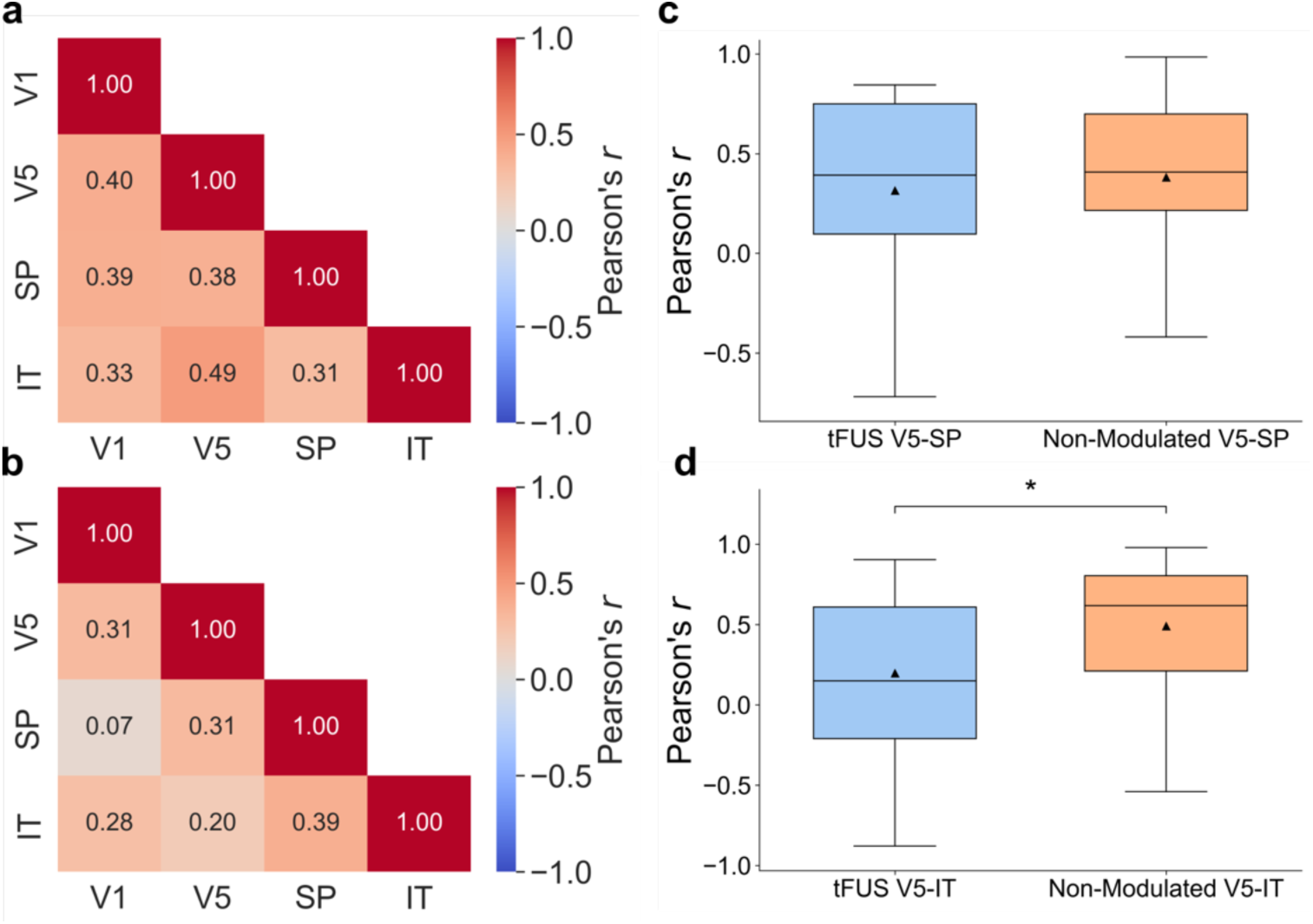
Pearson correlation coefficient (r) analysis of V1, V5, superior parietal lobe (SP), and inferior temporal gyrus (IT) connections for a) non-modulated and b) tFUS conditions. The mean correlation coefficient for each subject was averaged together, and indicate moderate correlation between V5-SP for both conditions. c-d) Comparisons of V5-SP and V5-IT correlations were made across conditions using a one-tailed Wilcoxon ranked sum test. d) No significant difference (*p* > 0.05) was found between the V5-SP correlation across conditions, indicating the pathway was preserved. The non-modulated condition also had moderate V5-IT correlation, but this was significantly (*p* < 0.05) dissociated in the tFUS condition (d).

### A natural language processing-based shared control algorithm further improves the BCI speller usability

Initial testing of a Shared Control autoRegressive Integrated Bayesian Estimator (SCRIBE; Sup. Fig. S1, Sup. Vid. S1, Sup. Vid. S2) suggests a shared control word-correction program can be used to further improve the software component of BCI spellers, especially for naïve human subjects, even when the typed accuracy is as low as 50%. While Bayesian autocorrection using word frequency is not a new concept for BCI spellers^41^, our algorithm considers both word frequency as well as keyboard-specific Euclidean error of letters. Further, existing Bayesian autocorrection formulas have considered just the current word^41^, while the presented SCRIBE considers words in the context of a *sequence*, thus imposing additional linguistic meaning onto the output. The current iteration only considers up to two-word sequences (autoregressive order 1), but there is available data on Google N Gram to consider up to five-word sequences (autoregressive order 4).

### *Ex-vivo* measurements of the applied tFUS parameters

tFUS target pressure and derated intensities generated by the customized 128-element random array ultrasound transducer H275 were estimated by transcranial pressure readings through 3D needle hydrophone scanning behind a fully hydrated real human skull sample submerged in degassed water in a water tank. The following ultrasound parameters are calculated for the administered transcranial ultrasound stimulation: 0.2 MPa peak-to-peak pressure at target, 233 mW/cm^2^ I_SPTA.3_, and 0.389 W/cm^2^ I_SPPA.3_. While currently there are no FDA safety guidelines specifically for ultrasound neuromodulation, the estimated ultrasound energy are within safety guidelines for ultrasound diagnostic imaging (I_SPTA.3_ < 720 mW/cm^2^, I_SPPA.3_ < 190 W/cm^2^)^42^. Transcranial ultrasound simulations showed the focused cortical stimulation without the presence of standing waves (Sup. Fig. S2).

## 3. DISCUSSION

Our study demonstrates, for the first time, that low-intensity transcranial focused ultrasound neuromodulation can be used to enhance the performance of a visual motion based brain-computer interface through significant theta amplification of the dorsal visual processing pathway. 0.2 MPa tFUS was administered at 3 kHz pulse-repetition frequency to a subject’s left hemisphere V5 while they typed letters through a virtual keyboard presented on a screen with an mVEP BCI speller. Euclidean errors of this BCI speller in tFUS condition were significantly lower than those of non-modulated, decoupled-sham, and US-control conditions, and ESI source imaging of the ultrasound-targeted left hemisphere V5 parcellation indicates significantly amplified N200 theta power. This amplification is observed to be carried downstream through the dorsal visual processing pathway, as the significantly amplified N200 theta power for the tFUS condition is also apparent in the superior parietal lobe. Correlation analysis supports the V5-superior parietal lobe information flow is maintained from the non-modulated to the tFUS condition, while extraneous pathways are disconnected. This, contextualized with previous theta-band research^43–48^, indicates that V5-targeted tFUS’ mechanism of action is to increase visual motion feature-based awareness.

EEG sensor domain-level analysis was conducted with a nonparametric permutation cluster test to account for the multiple comparisons problem. While the tests did reveal significant spatial-temporal (*p* < 0.05) and spatial-time-frequency (*p* < 0.05) clusters among the conditions, it should be noted that significant clusters do not guarantee underlying significant differences for all time points within the cluster, only that the clusters on average are significantly different^49^. Another potential limitation of the sensor-domain level data is differing sensor locations among subjects. EEG caps were not centered with respect to subject landmarks, but rather were rotated on an individual basis to allow for maximal exposure of the subject’s V5 for the placement and coupling to the ultrasound transducer. This may have led to different relative locations of EEG electrodes for each subject. This limitation of the spatiotemporal cluster test and differences in electrode positions were mitigated by adopting EEG electrode digitization for EEG source analysis. Captured EEG electrode placements were used for subject-specific ESI computations of various brain parcellations. Lowering the multiple comparison dimensionality from multiple electrodes at various timepoints to a summed theta power over the N200 time window allowed for direct comparisons across conditions with Kruskal-Wallis and Wilcoxon ranked sum testing.

The EEG source analyses were implemented using the theta component of the N200. The time-frequency permutation cluster test (Fig. 2d), as well as previous studies^43,50–52^, have established the significance of the theta band in the N200 conglomerate. Analysis of the superior parietal lobe N200 theta power suggests that effects of tFUS on V5 may carry on downstream in the dorsal processing pathway. The brain source activities in the tFUS condition are significantly amplified compared to those in the non-modulated and decoupled-sham conditions, and there is no significant difference found between the decoupled-sham and non-modulated conditions. The tFUS amplification directly reflects what was found in the V5 analysis, which is as expected given the canonical dorsal pathway, which involves projection of V5 to the super parietal lobe^38,40^.

N200 theta power analysis of the inferior temporal lobe shows no significant difference among conditions, which aligns with the fact that V5 canonically does not project to the ventral pathway^38,39,53^. Subject-average whole-brain ESI visualizations (Sup. Fig. S3) also show elevated V1 area activity in the tFUS condition for the N200 time window. The significance of the V5-V1 projection in processing visual motion has previously been established, and connectivity analysis yielded a moderate correlation (*r* > 0.3; Fig. 5a-b) between the cortical regions, so it would follow that the strengthened V5 activity from tFUS may result in amplified V1 activity for the visual motion BCI paradigm. Source analysis of V5 also revealed a significant damping of the decoupled-sham compared to the non-modulated condition. This might be due to the ERP response being affected by attention^47^, and the beeping noise generated by the PRF of the decoupled-sham ultrasound may have distracted the subjects without the actual pressure waves delivered in the tFUS condition to counteract the effect. Future research should continue to work towards a better understanding of the ultrasound auditory confounds in such a neuromodulation-assisted human BCI, as well as engineering strategies to mitigate potential side effects therein^54,55^.

Theta-band correlation analysis revealed that the non-modulated V5 had connectivity with the inferior temporal gyrus (Fig. 5a), which was significantly attenuated during the tFUS condition (Fig. 5d). The dorsal pathway connectivity (V5-superior parietal lobe), on the other hand, was preserved across conditions (Fig. 5c). These results indicating tFUS’s ability to break connections are supported by a non-human primate fMRI study^56^. The mechanism of action behind this disconnection may be directly related to the observed increased theta power, as previous work suggests increased theta activity is tied to inhibitory processes^43–47^. Following this, it is possible that the tFUS-induced amplified theta power in V5 inhibited extraneous connectivity (such as that from V5-inferior temporal gyrus). This hypothesis is also supported by a previous tFUS study suggesting the mechanism of action is to increase activity of local interneurons^22^.

This theory may be further supported by the fact that theta-band activity is also associated with attention^48^. The significance of visual feature-based attention creating a cognitive spotlighting effect has been widely studied with regards to V4 and color, shape, and orientation^57^. Since V4’s primary function is associated with object recognition^58^, which would include color, shape, and orientation, and V5’s primary function is associated with visual motion perception^32,33^, it follows that V5 modulation may draw heightened cognitive awareness to visual motion based features. Our data, in the context of these past studies, suggest that tFUS’ mechanism of action is to enhance feature-based attention associated with the stimulated brain region through amplified theta activity.

A significant disconnection (*p* < 0.05) is also seen for the V1-superior parietal lobe pathway during the tFUS condition (Fig. 5a-b). This result supports the previous hypotheses, as N200 theta analysis indicated significantly amplified theta power in the superior parietal lobe (Fig. 3c). This increased theta power may have resulted in a disconnection of a non-relevant information carrying pathway for the task (i.e., superior parietal lobe to/from V1), similar to what it induces at the site of sonication (i.e., the disconnection of V5 and IT). In the context of feature-based attention, V5 stimulation was carried through into the superior parietal lobe, as evident from the theta power and correlation analyses. This may have focused the function of superior parietal lobe on visual motion-based features, and, by doing so, may lead to weakening other connectivity in and out of the superior parietal lobe, including those with V1.

Strangely, a moderate correlation between the superior parietal and the inferior temporal gyrus is maintained across conditions (Fig. 5a-b; *r* > 0.3; *p* > 0.05). The dorsal and ventral processing pathways are canonically separate, but these results suggest there may be some interconnections between these two regions. In this study, the V5-inferior temporal gyrus connection was reduced during the tFUS condition, and no theta power change was observed across conditions in the inferior temporal gyrus. As such, it follows that whatever connectivity the two regions have, it is likely unassociated with visual motion feature-based attention. Future work is warranted to further assess this connection.

Significant reduction in BCI Euclidean error, as well as significant amplification of N200 theta power in V5 and the superior parietal lobe, of V5-targeted tFUS compared to the US-Control condition provide evidence that effects of tFUS are target specific, rather than a result of ultrasound energy projected nonspecifically into the brain. Modulating the US-Control location yielded lowered N200 theta power in the inferior temporal gyrus area. It is possible that sonicating the control location led to suppression of that area, either directly or indirectly, but more future research is warranted on investigating tFUS effects on the ventral pathways.

tFUS neuromodulation was implemented concurrently during the BCI task, meaning that it modulated brain signals while the visual stimuli were presented. This is different from traditional non-invasive neuromodulation studies for BCI, which are usually used before or after the trials to either prepare the brain or consolidate learning^19–21^. The technique in this study represents a step towards more closely mimicking invasive electrical stimulation^59^.

One limitation of this study has to do with the way the N200 theta powers are calculated. The process involved taking the mean of the power within 4 to 8 Hz in the 100 to 250 ms post-stimulus time window. N200 peak times are commonly found between 150 and 250 ms post onset^60–63^, so to get the onset of the wave, an earlier time window was used. However, this does lead to potential overlap with the N100 ERP^31^ and the end of the window may overlap with the P300 ERP^64^. Visualization of the trial averages (Fig. 2b) indicate that this window, on average, corresponds to an N200 waveform. However, this may not be guaranteed for every trial. Future works may further explore automated ERP differentiation and characterization to best isolate one from another. However, given the large trial numbers, exclusion of statistical outliers, and equal likelihood of this issue across all trial conditions, it is likely that the results are a mostly accurate representation of the N200 theta power.

In conclusion, tFUS neuromodulation to human V5 can significantly decrease mVEP-based BCI spelling error. Concurrent neural electrophysiological recordings indicate that in the mVEP-based BCI, tFUS neuromodulation causes a significant amplification of theta power in the immediate V5-targeted area, as well as downstream in the dorsal vision processing pathway. Connectivity analysis and previous theta-band research provide critical evidence that this increased theta power may lead to heightened feature-based attention of visual motion stimuli.

## 4. METHODS

### Participants

This study was reviewed and approved by the Advarra Instructional Review Board. 21 healthy human subjects (11 male / 10 female; mean age: 24.4 ± 5.92) were recruited from around the university area in Pittsburgh. Interested subjects were safety screened for MRI eligibility and informed of potential risks of the study. Subjects who wished to continue gave voluntary informed consent in accordance with the World Medical Association’s Declaration of Helsinki. A subject’s data were excluded from analysis if their non-modulated BCI classifier was random chance.

### Transcranial ultrasound characterization

#### Ex-vivo transcranial measurements

A 128-element random array ultrasound transducer H275 with 700 kHz fundamental frequency (Verasonics; Kirkland, Washington, USA) was characterized in a free-water tank through a hydrated human skull fragment (Fig. 6d). A hydrophone was placed on one side of the skull, opposite the ultrasound transducer. The hydrophone was moved with a stepper motor to capture the transcranial pressure field in three dimensions. Estimated target pressure, derated spatial peak pulse average intensity (I_SPPA.3_), and derated spatial peak temporal average intensity (I_SPTA.3_), values were calculated to ensure the safety in compliance with the current available FDA guidelines^42^.

**Figure 6.**
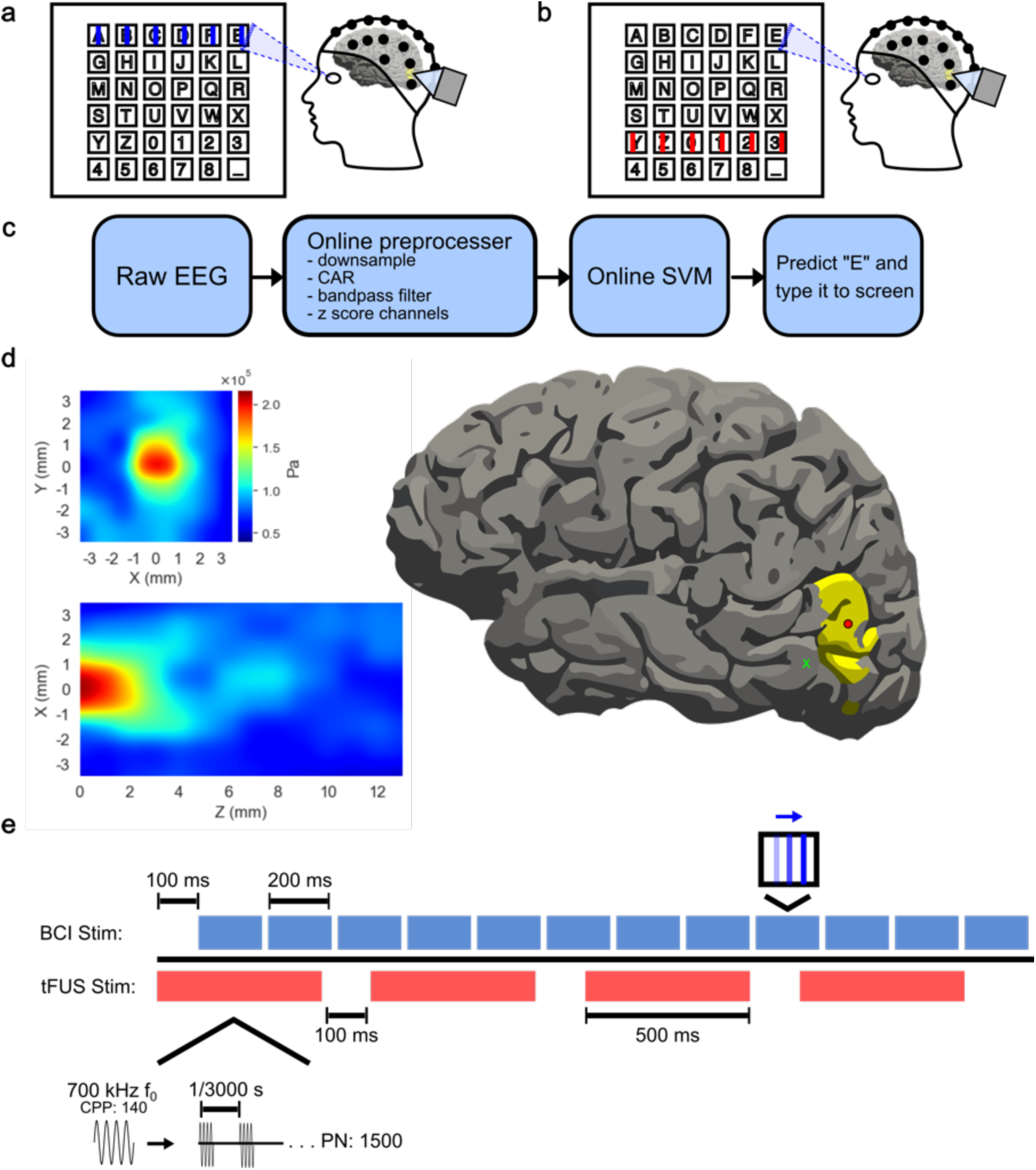
Experimental Paradigm for mVEP Speller. a) Example of “on-target” visual stimuli. The lines flash across the row or column of the letter the subject is looking at. b) Example of “off-target” visual stimuli. The lines flash on a row or column not associated with the letter of interest. c) The online BCI decoder pipeline. The raw EEG is down sampled to 100 Hz, bandpass filtered between 1 and 20 Hz, common average referenced, and z-scored across channels. The processed data is fed to an SVM classifier, which predicts the letter of intent based on neural activity for on and off target stimuli. d) (left) The 128 channel 700 kHz f_0_ tFUS pressure wave profile measured through a skull fragment in a water tank. (right) A visualization of the profiled 2 mm focal point scaled to a 93 mm brain and centered on V5 (red circle), as well as at an example US-Control area 1.41 cm away (green *x*). e) A typical trial epoch consists of 12 row/column 200 ms line flashes and four 500-ms tFUS sonications. The first sonication begins 100 ms prior to the first BCI stimulation.

#### In-silico transcranial simulations

Computer simulations of the random array ultrasound transducer were performed using Sim4Life (Zurich Med Tech; Zurich, Switzerland). The transducer was custom modeled and coded. The program’s standard head model with bone, tissue, and internal air pockets was used. The transducer was placed to roughly target the model’s visual cortex. The simulation considered 140 periods and 0.1 MPa transmission pressure from each element and was solved using the linear pressure wave equation (LAPWE) model.

### Neuronavigation

Each participant underwent a structure MRI, and the acquired brain images were further segmented into brain regions using FreeSurfer^35,36^. The participant’s MRI files and specific RAS (Right, Anterior, Superior) coordinates of the center of the segmented left hemisphere V5 were plugged into the Localite TMS navigator^65^. The physical size and orientation of H275 were calibrated to the navigation system and aimed at the participant’s left V5.

### Online mVEP BCI speller

Each participant’s head size was measured and was fit with an EEG cap. Incisions were made to the EEG cap to expose the V5 area, which allowed for direct interfacing between the scalp and the ultrasound transducer. Electrode placement relative to subject landmarks were captured for source imaging analysis. The electrode impedances were kept below 10 kΩ using conductive electrolyte gel and applied using cotton swabs.

The mVEP BCI speller was made in PsychoPy^66^ and designed based on previously published mVEP BCI literature^15^. Subjects were presented on screen with a six-by-six virtual keyboard. They were instructed to stare at the specific key they wished to type as lines flashed to the right across each row and column of the keyboard. One scan epoch consisted of lines flashing across each row and column once. For future analysis, “on-target” will refer to the lines that flashed across the row and column of letter the subject was looking at (Fig. 6a). “Off-target” will refer to the rest of the lines in the epoch (Fig. 6b). Depending on the subject, tFUS was pulsed 1, 2, or 4 times during each scanning epoch. The first sonication began starting 100 ms before the visual stimulus onset, and any additional doses were spaced equidistant over the remaining duration of the epoch.

A model-training session was conducted where subjects were instructed to type the descending diagonal (*AHOV2_*). The model was created offline using Scikit-Learn’s^67^ Support Vector Machine to predict the subject’s intended letter based on timing of differing neural activity of posterior electrodes *P2, P1, P3, P4, P5, P6, P7, P8, PO3, PO4, PO7, PO8, O1, O2, TP7,* and *TP8.* In five cases, additional electrodes *CP1, CP2, CP3, CP4, C1, C3, C2, C4, FC1, FC3, FC2, FC4* were also used. The same electrode set was used for all conditions for a subject. A trial consisted of data from stimulus onset to 500 ms after. Data were decimated to 60 Hz, bandpass filtered from 1 to 20 Hz, and z-scored with respect to each channel (Fig. 6c).

For the online testing session, 12 subjects were asked to type 14 letters (*CARNEGIEMELLON*) with active SCRIBE. The other nine subjects were asked to type 15 letters (*CARNEGIE_MELLON*) without SCRIBE. Online EEG data were processed and fed to the classifier once per scan epoch. For each epoch, the classifier predicted the maximally probable row and column index of the subject’s gaze. If there were multiple epoch scans per letter, the classifier would average the probabilities over the epochs.

This process was done for four conditions in a randomized order: “non-modulated” (inactive tFUS), “decoupled-sham” tFUS (decoupled, but active, tFUS to account for audio-induced confounds), “US-control” (active tFUS targeted to a control brain location located by more than 1-cm away), and V5-targeted “tFUS” (estimated peak-to-peak pressure: 0.2 MPa, approximate focal beam diameter: 2 mm, approximate focal beam length: 2 mm (Fig. 6d), pulse repetition frequency: 3 kHz, pulse duration: 200 µs, sonication duration: 500 ms (Fig. 6e)). In some cases, time did not allow for testing all four conditions and subjects were unable to be scheduled back. The partial data for these sessions were included in analysis if they did not meet the other exclusion criteria.

### BCI performance analysis

BCI performance was evaluated for each condition’s online testing session based on the Euclidean distance error (EE) from the subject’s intended letter to the BCI classifier’s output. The output for each letter (i.e., *C, A,* etc) was considered as one separate trial, and all the trials for all the subjects were pooled together. The Euclidean distance was converted to a percent error by normalizing it against the maximum possible distance (diagonal distance of the grid: 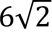). The results were also normalized with respect to subject-specific ultrasound dosage and the square root of the number of scan epochs the BCI averaged over (Equation 1). For the non-modulated condition, the final term was omitted. Euclidean errors were compared across conditions using a Kruskal-Wallis test and a two-tailed Wilcoxon ranked sum test. Bonferroni-Holm *p* adjustment was applied to reduce likelihood of family-wise errors. Euclidean errors were considered significantly difference if *p*_adjusted_ < α = 0.05.

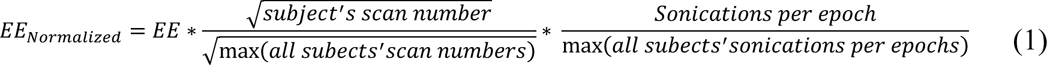

### Offline Neural data analysis

Data analysis was performed in Python^68^ using MNE^37^ and SciPy^69^ packages for EEG processing and statistical testing. Since testing error was nonzero, it was not always clear whether the subjects were looking at the keys they were intended to. Therefore, to maximize the likelihood of properly matching the visual stimuli to “on-target” / “off-target” labels, electrophysiological signal analysis was performed on the EEG data collected during the BCI training session. Each scanning epoch was considered as one trial.

#### EEG preprocessing

EEG were bandpass filtered between 1 to 40 Hz, common average referenced, and down-sampled to 100 Hz. Independent component analysis (ICA) was performed to remove eyeblink-related artifacts using MNE Python’s automatic EOG detection, considering FP1 as an EOG channel with a z-threshold of 3.0^70^. Bad epochs were rejected and sensors were cleaned using the autoreject package^71,72^, and persisting eyeblink ICA bases were rejected with an additional run of MNE’s automatic EOG detection using both FP1 and FP2 and a z-threshold of 3.5.

#### EEG sensor spatiotemporal analysis

Data from all subjects were pooled into one dataset per condition and z-scored with respect to the 200 ms baseline activity before stimulus onset. Sensor-domain level data for the tFUS condition were analyzed against decoupled-sham and non-modulated conditions using two MNE Python nonparametric permutation cluster tests^73^. For this, an F-test was conducted across all time and space to compare the two tested conditions, and significantly different (α *=* 0.05), points that were adjacent in time and space were clustered together. Then, data across conditions were shuffled into random permutations of the original conditions. An F-test was run for the values at all time points and electrodes. Significant (α *=* 0.05) adjacent time points and electrodes were clustered together. The number of datapoints in the largest cluster was stored and used to generate a cluster-size distribution. The cluster-forming size threshold was calculated as the 95^th^ percentile (*p* < 0.05) from the distribution. Clusters from the original data comparison that were larger than the threshold size were deemed significant spatiotemporal clusters. This test used 1,000 permutations. Significant clusters that did not include posterior electrodes or took place prior to stimulus onset were omitted.

#### EEG sensor time-frequency analysis

A permutation cluster test was also implemented for time-frequency responses. Preprocessed EEG data were pooled into one dataset per condition and computed with time-frequency representations using Morlet wavelets for frequencies between 3 and 20 Hz at an interval of 3 Hz. The transformed time-frequency data were then baseline corrected by subtracting the mean value of its 200 ms pre-stimulus baseline. This test considered differences across all three conditions. It ran the same as the spatiotemporal test, except for that is also considered frequency decomposition. Significant (α *=* 0.05) adjacent time-frequency-space data were clustered together both for the original datasets and for random permutations of the data. The cluster-forming size threshold was determined based on the probability distribution cut off for the F statistic > 10. Clusters from the original data comparison that had a size greater the cut-off were deemed significant. The test ran for 1,000 permutations. Only posterior left hemisphere electrodes (*O1*, *PO3*, *P1*, *P3*, *P5*, *P7*, *TP7*, and *PO7*) were considered for this test. The same exclusion criteria were used as the spatiotemporal analysis.

#### EEG source imaging

Time series current density data for the left hemisphere V5 region was extracted from the preprocessed EEG data using MNE Python in the FreeSurfer^35,36^ segmented region (Brodmann Area (*BA_exvivo*) atlas’ *MT_exvivo-lh* label). MRI and EEG digitization data were aligned using MNE Python’s automatic co-registration function^74^. The source imaging problem was solved with Minimum Norm Error method^75,76^. In five cases, a subject’s MRI had mesh errors that did not allow for source imaging. In these scenarios, MNE Python’s base head model was used instead of the subject specific MRI model. The source imaged time series were bandpass filtered from 1 to 40 Hz and z-scored with respect to each trial’s 200 ms pre-stimulus baseline.

Source imaged time series were transformed into time-frequency power responses using Morlet waveforms^77^. The N200 theta power was extracted by taking the 4 to 8 Hz frequency power response in the 100 to 250 ms post-stimulus window. Outlier trials were rejected by the interquartile range test. Shapiro and F-tests were used to evaluate data normality and equality of variances across conditions, respectively. As a result, theta powers were compared across conditions using a Kruskal-Wallis test and a two-tailed Wilcoxon ranked sum test with Bonferroni-Holm *p*-adjustment (α *=* 0.05).

This process was repeated with the Desikan-Killiany (*aparc*) atlas’ parcellations for the left hemisphere superior parietal lobe (*superiorparietal-lh*) and the left hemisphere inferior temporal lobe (*inferiortemporal-lh*) to examine possible downstream effects in visual processing pathways.

#### Connectivity analysis

Cortical functional connectivity was assessed using Pearson’s R correlation coefficient. The average current density time series at V1, V5, the superior parietal lobe, and the inferior temporal gyrus were calculated for each subject, bandpass filtered 1 to 40 Hz, z-scored with respect to the 200 ms baseline, and then filtered from 4 to 8 Hz to extract the theta band component. A correlation coefficient was calculated for each connection from stimulus onset to 500 ms afterwards in the tFUS and non-modulated condition. Changes in the V5-IT and V5-SP correlations were tested with a one-tailed Wilcoxon ranked sum test.

### <u>S</u>hared <u>C</u>ontrol auto<u>R</u>egressive <u>I</u>ntegrated <u>B</u>ayesian <u>E</u>stimator (SCRIBE)

In order to further improve the mVEP speller classifier, a custom autocorrect program was created. This program was based on the principles of Bayesian inference, such that:

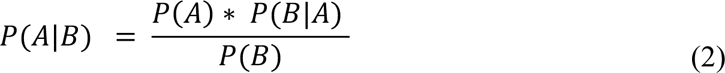

in which *A* is corrected word, and *B* comprises the typed letters. The probability of the *corrected word* was estimated by the square-root log-frequency of the word’s occurrence based on Google N Gram’s 2018 database^78^. The probability of the typed letters given the corrected word was defined as the Euclidean error between the two. The probability of typed letters occurring would be the same for all possible corrected words, so the denominator was not calculated. Thus:

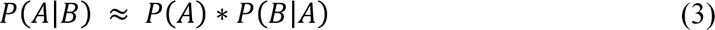

in which *P(B|A)* is the probability of the typed letters given the corrected word, and *P(A)* is the probability of the corrected word. This can be taken a step further, as language follows certain patterns, and some words become more probable following others. If the current word is following at least one more word, the formula changes to incorporate the past words:

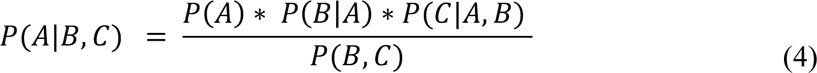

in which *C* comprises the previous word(s). We can make the assumption that the previous word is independent of the current letters, leading to:

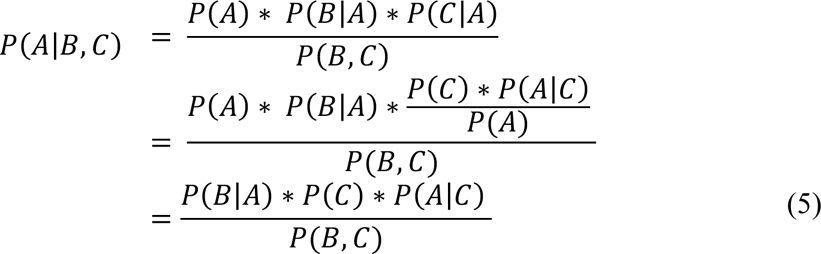

For every possible corrected word, the probability of currently typed letters occurring (*P(B*)), the probability of the previous word(s) occurring (*P(C)*), and the joint probability of the two (*P(B, C)*) are the same. Therefore, to increase computation efficiency, these terms can be ignored in the calculation. The new equation becomes:

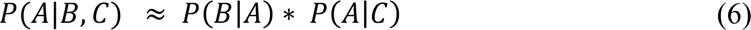

in which *P(B|A)* is the probability of the typed letters given the corrected word, and *P(A|C)* is the probability of the corrected word given previous word(s). The probability of the *corrected word* occurring given *previous word(s)* was estimated by the scaled log-frequency of the singular words and word sequences from data in Google N Gram’s 2018 multiword database.

This was tested during the online BCI session as a greedy search (Sup. Vid. S1, Sup. Vid. S2). Probabilities were calculated for all words in the Scrabble dictionary, as well as “Carnegie” and “Mellon.” Three possible options were given to the subject at the end of every word. These options were the original letters typed and the next two most probable real words. This means that if the user happened to type a real word, they would be provided with the three most probable word options.

### Data availability

The data are presented in the paper and supplementary materials. Additional data will be made public through a data repository upon paper acceptance.

### Code availability

Computer codes will be made public via GitHub when the paper is published.

## Acknowledgments

This work was supported in part by NIH grants R01NS124564, R01AT009263, U18EB029354, T32EB029365, and R01NS096761. The authors would also like to thank Jenn Shanahan for her assistance in EEG capping, and Yunruo Ni for her assistance in EEG capping and 3D printing the tFUS transducer holder.

## Author Contributions

**Conceptualization:** J.K., K.Y., and B.H. **Methodology:** J.K., K.Y., and C.L. **Data collection:** J.K. and K.Y. **Formal analysis:** J.K. **Investigation:** J.K., K.Y., and B.H. **Writing – Original draft:** J.K. **Writing – Reviewing and editing:** J.K., K.Y. and B.H. **Supervision:** K.Y. and B.H.

