## Supplemental Figures S1-S3 for "Transcranial Focused Ultrasound to V5 Enhances Human Visual Motion Brain-Computer Interface by Modulating Feature-Based Attention"

**of**

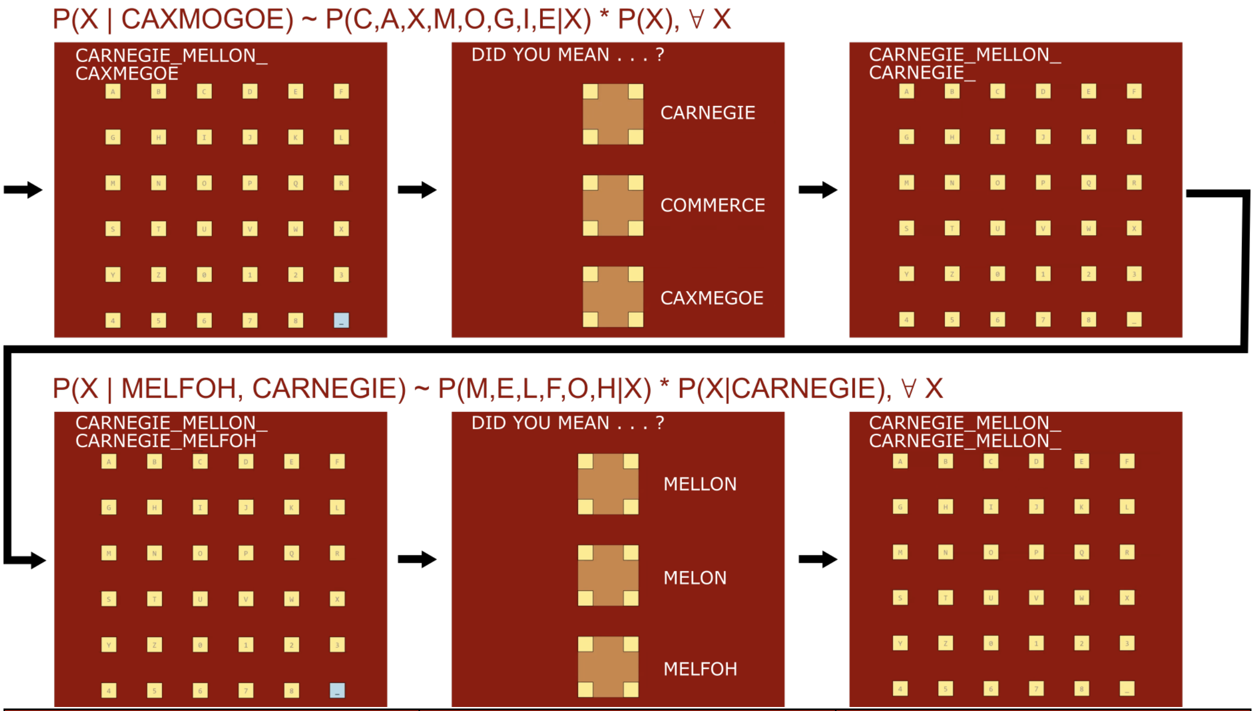


| Typed Letters | Original Typed Accuracy | Online SCRIBE Result |
|---|---|---|
| CARNEGCE MAJLMN | 71.4% | CARNEGIE MELLON |
| CARNAGIE MBLLON | 85.7% | CARNEGIE MELLON |
| CARNEGGW MEIL0H | 64.3% | CARNEGIE MELLON |
| CARNWLID MELXOM | 71.4% | CARNEGIE MELLON |
| CAQOEG0E KE3WOC | 50.0% | CARNEGIE MELLON |
| CA_NEAOE MGRLTN | 57.1% | CARNEGIE MELLON |
| CAXMEGOE MELFOH | 64.3% | CARNEGIE MELLON |
| CALNEAIE QELLON | 78.6% | CARNEGIE MELLON |
| CARNDAIK AELLOH | 71.4% | CARNEGIE MELLON |
| CARN2GLE MECLON | 78.6% | CARNEGIE MELLON |

**Supplemental Figure S1.** Shared Control autoRegressive Integrated Bayesian Estimator (SCRIBE) can improve BCI speller performance by applying linguistic rules**.** (top) Diagram of SCRIBE algorithm based on an online test (Sup. Vid. 1 bottom right). At the end of every word, a greedy search is performed based on the typed letters and previous words to determine the most probable corrections (Equation 6). If there is no previous word, the probably of the singular word occurring is considered instead. The two most probable words and the originally typed letters are presented as correction options. When the user makes a selection, the previous word is updated to reflect the choice. (bottom) Selected examples of online SCRIBE results. Even when the user's actual accuracy is 50%, the words are still able to be corrected by considering a combination of Euclidean errors and linguistic patterns.

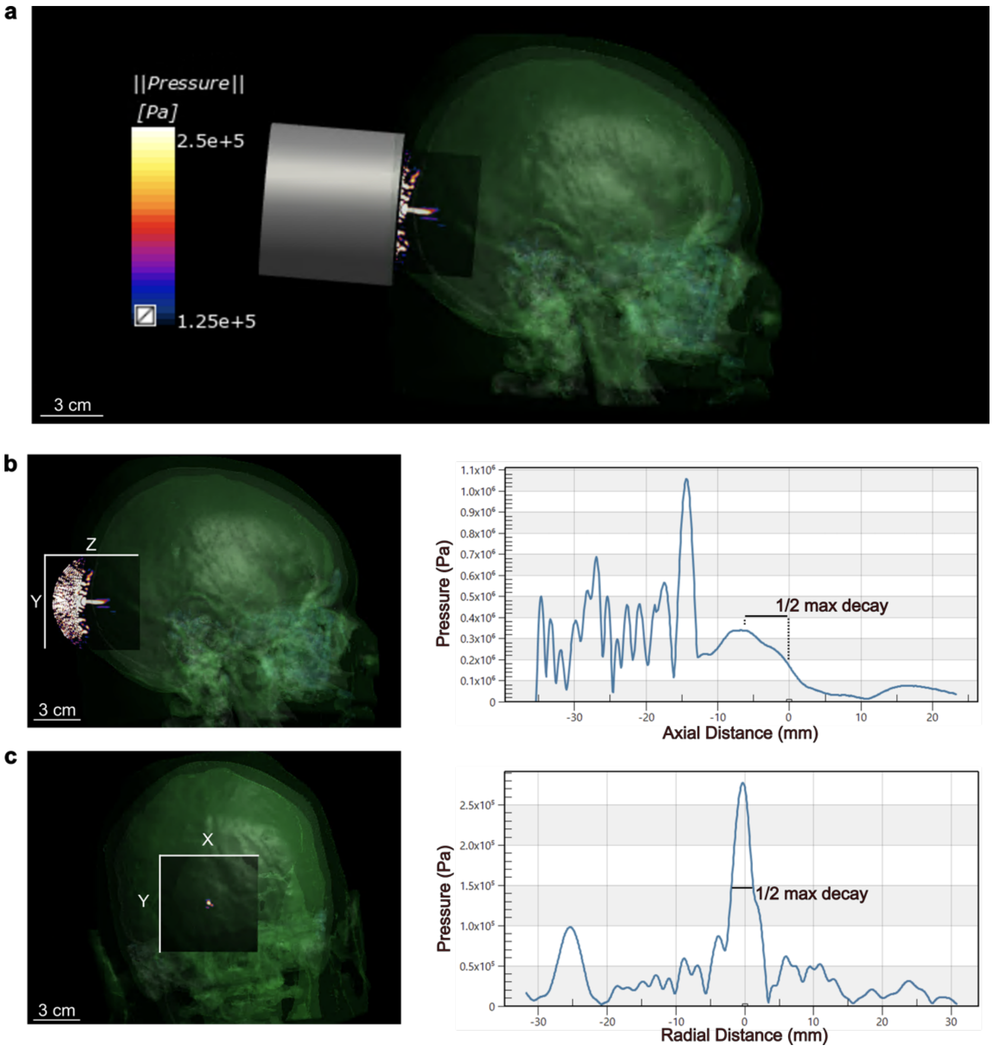


**Supplemental Figure S2.** a) 128-element random array 700 kHz ultrasound transcranial simulation (140 periods) with Sim4Life on a human head (using CT images). b) The simulation results for intracranial pressure in the YZ plane. The approximate transcranial ultrasound beam length is 6.76 mm (the full width at half maximum, FWHM). c) The simulation results in the XY plane. The approximate FWHM after the skull is 3.67 mm. The simulation does not result in standing waves forming inside the skull cavity of the head model. The focus of tFUS is localized at the visual cortical brain, and does not reach deep brain structures.

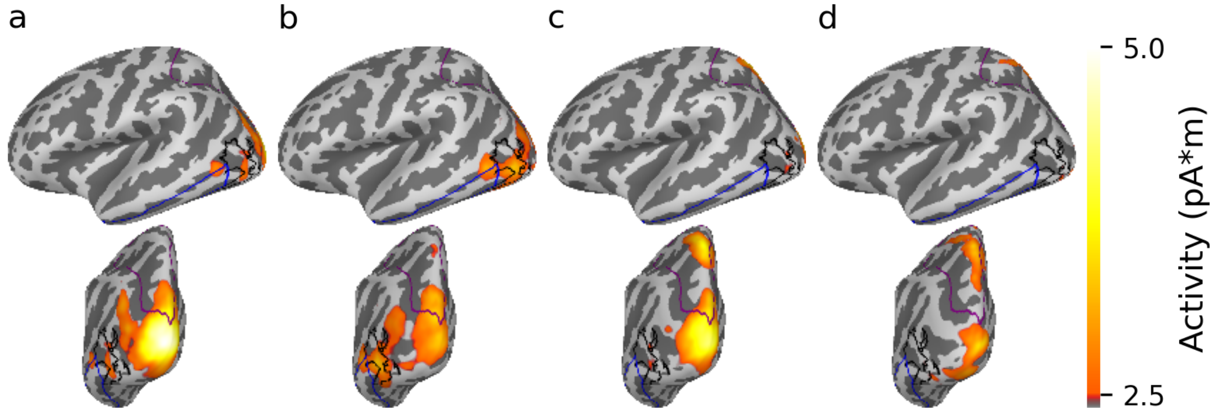


**Supplemental Figure S3.** EEG source imaging from all subjects projected onto an average brain model of 40 MRI scans (FreeSurfer's *FSAverage*[79]). EEG data for each subject was first fit to their head model and then source morphed to the *FSAverage* model. The activity across all subjects on the *FSAverage* model was averaged. Labeled regions for V5, the superior parietal lobe, and the interior temporal gyrus are outlined in black, purple, and blue, respectively. The data were bandpass filtered between 1 and 40 Hz Data and baseline corrected by subtracting the mean of the 200 ms prestimulus window. Data were further averaged over the 100 to 250 ms post stimulus window. Conditions: a) tFUS, b) Non-Modulated, c) Decoupled-Sham, and d) US-Control. The tFUS condition had the highest single-voxel amplitude (4.95 pA*m; non-modulated: 3.45 pA*m, Decoupled-Sham: 4.39 pA*m, US-Control: 4.39 pA*m).
